# Emergence of plasmid-mediated RND-type efflux pump gene cluster *tmexCD-toprJ* in *Shewanella xiamenensis* in a water environment

**DOI:** 10.1101/2022.09.15.508154

**Authors:** Trung Duc Dao, Taichiro Takemura, Ikuro Kasuga, Aki Hirabayashi, Nguyen Thi Nga, Pham Hong Quynh Anh, Nguyen Dong Tu, Le Thi Trang, Hoang Huy Tran, Keigo Shibayama, Futoshi Hasebe, Masato Suzuki

## Abstract

The emergence of the mobile resistance-nodulation-division (RND)-type efflux pump *tmexCD-toprJ* gene cluster that confers multidrug resistance (MDR), including tigecycline resistance, in gram-negative bacteria poses a global public health threat. However, the spread of such clinically important antimicrobial resistance genes (ARGs) in the natural environment has not yet been well investigated. In this study, we investigated MDR aquatic bacteria in Vietnam. A carbapenem- and tigecycline-resistant *Shewanella xiamenensis* isolate NUITM-VS2 was obtained from urban drainage in Hanoi, Vietnam, in October 2021. *S. xiamenensis* NUITM-VS2 showed resistance to most antimicrobials tested, including tigecycline, tetracyclines, carbapenems, cephalosporins, fluoroquinolone, and aminoglycosides. Whole-genome analysis was performed by long- and short-read sequencing, resulting in the complete genome sequence consisting of one chromosome and five plasmid sequences. ARGs and plasmid replicons in the genome were detected using ResFinder with the custom ARG database, including all known tigecycline resistance genes, and PlasmidFinder, respectively. A 152.2-kb IncC plasmid, pNUITM-VS2_2, co-carried two mobile tigecycline resistance genes, *tet*(X4) and *tmexC3.2D3.2-toprJ1*. In addition, a 24.8-kb untypeable plasmid, pNUITM-VS2_4, carried the carbapenemase gene *bla*_NDM-1_. pNUITM-VS2_2 was transferred to *Escherichia coli* by conjugation, which simultaneously conferred high-level resistance against many antimicrobials, including tigecycline. To the best of our knowledge, this is the first report of the detection of the mobile RND-type efflux pump gene cluster *tmexCD-toprJ* in *Shewanella* species. Our results provide genetic evidence of the complexity of the dynamics of clinically important ARGs among aquatic bacteria, which could be important reservoirs for ARGs in the natural environment.

## Introduction

Tigecycline is a semi-synthetic antimicrobial agent, which was the first drug of the glycylcycline class developed for the last-line treatment of infections caused by multidrug-resistant (MDR) bacterial pathogens, especially carbapenem-resistant *Enterobacterales.* However, due to the widespread use of this antimicrobial in clinical settings, novel antimicrobial resistance genes (ARGs), including the mobile tigecycline-inactivation enzyme gene *tet*(X) and the resistance-nodulation-division (RND)-type efflux pump gene cluster *tmexCD-toprJ*, both conferring high-level tigecycline resistance, have rapidly been disseminated between diverse gram-negative bacteria (1). This not only poses an economic burden to the healthcare system but also increases mortality risks in patients by disabling the availability of last-resort antimicrobial agents, such as tigecycline.

*Shewanella* species are gram-negative anaerobes found in marine and freshwater environments, which belong to the family Shewanellaceae within the class Gammaproteobacteria. *Shewanella* are natural reservoirs of *bla*_OXA-48_-like carbapenemase genes, such as *bla*_OXA-48_ and *bla*_OXA-181_, encoded on the chromosomes (2, 3). In 2022, we reported a *Shewanella xiamenensis* isolate obtained from a water environment in Vietnam, which harbored *bla*_OXA-48_ and *tet*(X4) on the chromosome (4).

In 2021, we reported a *Klebsiella aerogenes* isolate harboring a self-transferable incompatibility group C (IncC)–IncX3 hybrid plasmid co-carrying *bla*_NDM-4_, *tet*(X4), and *tmexCD3-toprJ1* [initially named *tmexCD3-toprJ3* (5) and later renamed (6)] from a water environment in Vietnam (7). Although the *Flavobacteriaceae* family is estimated as the natural reservoir of *tet(X)* genes (8), limited information is available on the distribution of *tmexCD-toprJ* in the natural environment.

In this study, we report an *S. xiamenensis* isolate obtained from the aquatic environment in Vietnam harboring *tmexCD-toprJ,* representing the first such report for *Shewanella* species. The isolate showed carbapenem and tigecycline resistance, and harbored *bla*_OXA-181_ on the chromosome, *bla*N_DM-1_ on a 24.8-kb untypeable plasmid, and *tmexC3.2D3.2-toprJ1* together with *tet*(X4) on a self-transferable 152.2-kb IncC plasmid.

## Results and discussion

An environmental carbapenem- and tigecycline-resistant bacterial isolate, NUITM-VS2, was identified as *Shewallena* species using MALDI Biotyper. The minimum inhibitory concentrations (MICs) of tigecycline, tetracyclines, carbapenems, cephalosporins, and gentamycin were all above 128 mg/L (R); and those of ciprofloxacin, tobramycin, and streptomycin were 4, 16, and 64 mg/L (R), respectively; whereas that of amikacin was 0.5 mg/L (S) (Table 1).

**Table 1.** MICs of antimicrobials against *S. xiamenensis* NUITM-VS2 and the transconjugant of *E. coli* J53. Abbreviations: TIG, tigecycline; MIN, minocycline; DOX, doxycycline; TET, tetracycline; IPM, imipenem; MEM, meropenem; CTX, cefotaxime; CAZ, ceftazidime; CIP, ciprofloxacin; AMK, amikacin; GEN, gentamicin; TOB, tobramycin; STR, streptomycin.

| Strain | MIC (mg/L) |  |  |  |  |  |  |  |  |  |  |  |  |
| --- | --- | --- | --- | --- | --- | --- | --- | --- | --- | --- | --- | --- | --- |
|  | TIG | MIN | DOX | TET | IPM | MEM | CTX | CAZ | CIP | AMK | GEN | TOB | STR |
| *S. xiamenensis*<br>NUITM-VS2 | >128 | >128 | >128 | >128 | >128 | >128 | >128 | >128 | 4 | 0.5 | >128 | 16 | 64 |
| *E. coli*<br>J53/pNUITM-VS2_2 | 128 | 128 | 128 | 128 | 0.25 | 0.016 | 0.032 | 0.25 | 0.016 | 0.5 | 0.5 | 0.25 | 0.5 |
| *E. coli*<br>J53 | 0.25 | 2 | 4 | 1 | 0.25 | 0.016 | 0.032 | 0.25 | 0.016 | 0.5 | 0.5 | 0.25 | 0.5 |
| *E. coli*<br>ATCC 25922 | 0.25 | 1 | 2 | 1 | 0.25 | 0.016 | 0.064 | 0.5 | 0.008 | 1 | 0.5 | 0.5 | 0.5 |

Whole-genome analysis using the Illumina and Oxford Nanopore Technologies (ONT) sequencing platforms resulted in the complete genome sequence consisting of one chromosome and five plasmids, designated pNUITM-VS2_1, pNUITM-VS2_2, pNUITM-VS2_3, pNUITM-VS2_4, and pNUITM-VS2_5 (accession nos. AP026732, AP026733,AP026734, AP026735, AP026736, and AP026737), respectively. Genome annotation and average nucleotide identity (ANI) analyses revealed that the 4.9-Mb chromosome with a 46.4% GC content contains 4,100 coding sequences (CDSs), in which *bla*_OXA-181_ was detected by ResFinder, showed 97.3% identity to the genome sequence of *S. xiamenensis* JCM 16212^T^ (accession no. BMOP01000000).

The 152.2-kb pNUITM-VS2_2 plasmid of *S. xiamenensis* NUITM-VS2 contained a replicon classified to IncC and co-carried two mobile tigecycline resistance genes, *tet*(X4) and a variant of *tmexCD3-toprJ1*, whose sequence was a perfect match to that reported in the *K. aerogenes* pNUITM-VK5_mdr plasmid (accession no. LC633285) isolated from the same environmental location as NUITM-VS2 in Hanoi, Vietnam, in February 2021 (7) (Fig. 1). The CDSs of *tmexCD-toprJ* in pNUITM-VS2_2 and pNUITM-VK5_mdr showed high identity to *tmexCD3-toprJ1* of *Proteus mirabilis* RGF134-1 (accession no. CP066833) (5) isolated from a pig in China in 2019. The identity for *tmexC3* was 99.7% (1,161/1,164 nt), with three amino acid substitutions (Q187H, T256M, and A386T). For *tmexD3,* the identity was 99.9% (3,133/3,135 nt), including two amino acid substitutions (V610L and L611F). For *toprJ1,* the identity was 100% (1,434/1,434 nt). Therefore, we named the variant identified in this and previous studies *tmexC3.2D3.2-toprJ1*, consisting of *tmexC3.2, tmexD3.2,* and *toprJ1.*

**Fig. 1.**
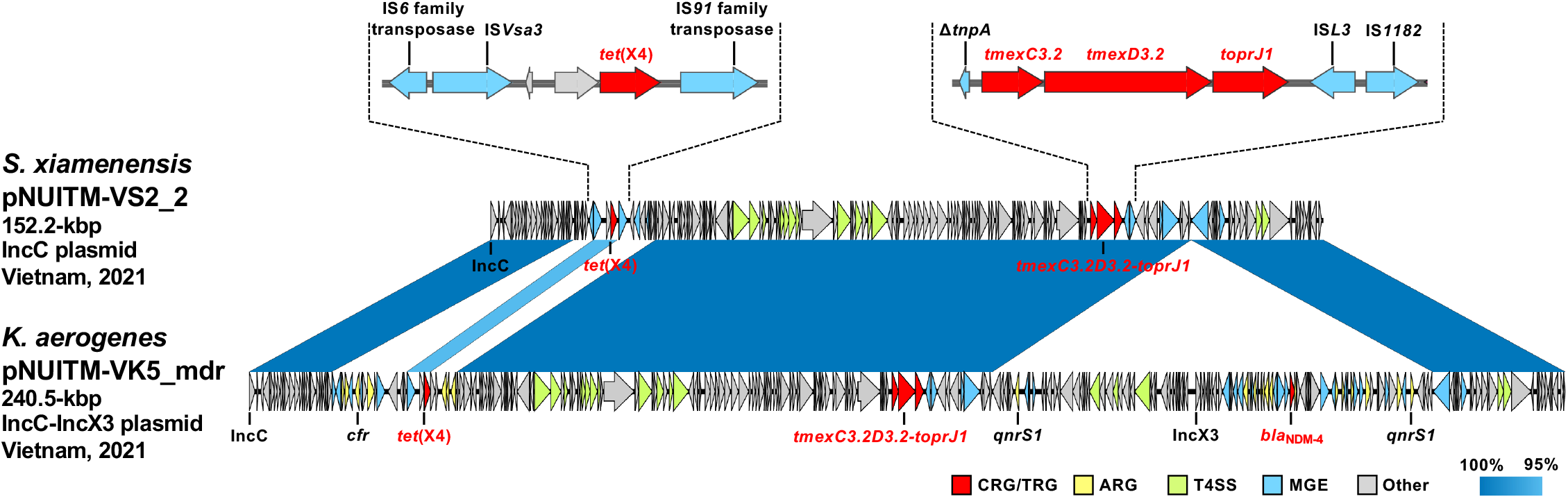
Linear comparison of *tet*(X4) and *tmexC3.2D3.2-toprJ1–carrying* plasmids *S. xiamenensis* pNUITM-VS2_2 and *K. aerogenes* pNUITM-VK5_mdr isolated in Vietnam in 2021. Red, yellow, blue, green, and gray indicate carbapenem and tetracycline resistance genes (CRG/TRG), other AMR genes (ARG), mobile gene elements (MGE), type IV secretion system (T4SS)-associated genes involved in conjugation, and other coding sequences (Other), respectively. The blue color in comparison of sequences indicates % of identity.

Moreover, the carbapenemase gene *bla*_NDM-1_, fluoroquinolone resistance gene *qnrVC6,* and aminoglycoside resistance genes *aac(3)-IId* and *aac(6’)-IIa* were encoded in another 24.8-kb untypeable plasmid, pNUITM-VS2_4, that is highly identical to the *S. xiamenensis* 26.8-kb untypeable plasmid pSx1 (100% identity with 96% of the region of accession no. CP013115) isolated from the environment in Algeria in 2012 (9). pSx1 was previously classified as an IncP-2 plasmid (10), which was subsequently found to be a misidentification (11). The replicon type of pNUITM-VS2_4 and pSx1 could not be identified by PlasmidFinder; thus, this plasmid type is presumed to be unique to *Shewanella* species.

On pNUITM-VS2_2 in *S. xiamenensis* NUITM-VS2, *tet*(X4) was flanked by two mobile genetic elements, IS*Vsa3* and *IS91* family transposase, while *tmexC3.2D3.2-toprJ1* was flanked by *ΔtnpA* and IS*Ł3*. Interestingly, pNUITM-VS2_2 was almost identical to the IncC backbone of the *K. aerogenes* IncC–IncX3 hybrid plasmid pNUITM-VK5_mdr (100% identity with 38% of the region of accession no. LC633285) (7) (Fig. 1). The rest of the IncX3 backbone of pNUITM-VK5_mdr carried additional ARGs, such as *bla*_NDM-4_ and *qnrS1,* conferring resistance to carbapenem and fluoroquinolone, respectively, suggesting that the MDR IncC–IncX3 hybrid plasmid pNUITM-VK5_mdr resulted from fusion of a tigecycline-resistance IncC plasmid such as pNUITM-VS2_2 with a carbapenem-resistance IncX3 plasmid.

A bacterial conjugation assay using *Escherichia coli* J53 as the recipient strain showed that *S. xiamenensis* NUITM-VS2 transferred pNUITM-VS2_2 to J53 at a frequency of 6.4 × 10^-5^ after overnight co-culture at 37°C. PCR confirmed that the transconjugant strain (J53/pNUITM-VS2_2) co-harbored *tet*(X4) and *tmexC3.2D3.2-toprJ1*, and was resistant to tigecycline and tetracyclines (Table 1).

## Conclusions

In conclusion, the complete genome sequence of the carbapenem- and tigecycline-resistant isolate *S. xiamenensis* NUITM-VS2 obtained from a water environment in Vietnam was characterized. This isolate co-carried two mobile tigecycline resistance genes, *tet*(X4) and *tmexC3.2D3.2-toprJ1*, in the self-transferable 152.2-kbp IncC plasmid. This plasmid corresponds to the IncC backbone of the IncC–IncX3 hybrid plasmid pNUITM-VK5_mdr identified in *K. aerogenes* in our previous report that was isolated from the same area (7), suggesting that they might share a common ancestor. Our results provide genetic evidence that clinically important ARGs have been disseminated though plasmids, such as broad-host-range IncC plasmids, and their fusion plasmids among environmental bacteria, such as *Shewanella* species, in aquatic environments.

## Materials and methods

### Bacterial isolation and antimicrobial susceptibility testing

Carbapenem- and tigecycline-resistant *S. xiamenensis* NUITM-VS2 was isolated from the Kim-Nguu River in Hanoi, Vietnam, in October 2021. Environmental water sample was collected and cultured using Luria-Bertani (LB) broth containing 4 mg/L of meropenem at 37°C overnight, and then further selected and isolated using CHROMagar COL-APSE (CHROMagar Microbiology) containing 4 mg/L of tigecycline. Bacterial species identification was performed using MALDI Biotyper (Bruker). Antimicrobial susceptibility testing using *Escherichia coli* ATCC 25922 as quality control was performed with agar dilution according to the CLSI 2020 guidelines. MIC breakpoints of antimicrobials (S: susceptible; I: intermediate; R: resistant) for other *non-Enterobacterales* were also determined according to the CLSI 2020 guidelines.

### Whole-genome sequencing and subsequent bioinformatics analysis

Whole-genome sequencing of *S. xiamenensis* NUITM-VS2 was performed using MiSeq (Illumina) with MiSeq Reagent Kit v2 (300-cycle) and MinION (Oxford Nanopore Technologies; ONT) with the R9.4.1 flow cell. The library for Illumina sequencing (paired-end, insert size of 300–800 bp) was prepared using the Nextera XT DNA Library Prep Kit, and the library for MinION sequencing was prepared using the Rapid Barcoding Kit (SQK-RBK004). ONT reads were base-called using Guppy v5.0.11, in the super-accuracy mode. Hybrid de novo assembly with both Illumina and ONT reads was performed using Unicycler v0.4.8.0 (https://github.com/rrwick/Unicycler) (12) with default parameters.

Genome annotation and average nucleotide identity (ANI) analyses were performed using the DFAST server (https://dfast.nig.ac.jp) (13). ARGs and plasmid replicons were detected using ResFinder v4.1 (https://bitbucket.org/genomicepidemiology/resfinder/) (14) with the custom ARG database, including all known tigecycline resistance genes, and PlasmidFinder v2.1 (https://bitbucket.org/genomicepidemiology/plasmidfinder/) (15) with the default parameters, respectively. Type IV secretion system (T4SS)-associated genes involved in conjugation were detected by CONJscan v1.0.5 (https://github.com/macsy-models/TXSS) with default parameters (16). Linear comparison of sequence alignment was performed using BLAST and visualized by Easyfig v.2.2.2 (http://mjsull.github.io/Easyfig/) (17).

### Bacterial conjugation assay

A bacterial conjugation assay was performed as follows. LB broth cultures of the donor *S. xiamenensis* NUITM-VS2 and the recipient azide-resistant *E. coli* J53 (ATCC BAA-2731, F^-^ *met pro* Azi^r^) were mixed in a 1:10 ratio, spotted onto MacConkey agar, and then incubated at 37°C overnight. Subsequently, the mixed cells, including transconjugants, were suspended in LB broth and then plated onto MacConkey agar containing 1 mg/L of tigecycline and 100mg/L of sodium azide after 10-fold serial dilution and incubated at 37°C overnight.

*tet*(X) and *tmexCD-toprJ* in transconjugants were detected by colony PCR using the following primer sets: tetX_F, CCCGAAAATCGWTTTGACAATCCTG; tetX_R, GTTTCTTCAACTTSCGTGTCGGTAAC; tmexC_F, TGGCGGGGATCGTGCTCAAGCGCAC; tmexC_R, CAGCGTGCCCTTGCKCTCGATATCG. The PCR amplification consisted of the initial denaturation for 3 min at 95°C, followed by 35 cycles of denaturation for 30 s at 95°C, annealing for 30 s at 54°C, and extension for 30 s at 72°C, and then the final extension for 5 min at 72°C. Genomic DNAs of *K. aerogenes* strain NUITM-VK5 co-harboring *tet*(X4) and *tmexC3.2D3.2-toprJ1* (7) was used as positive controls. Purified water was used as a negative control.

### Nucleotide sequence accession numbers

The complete genome sequence of *S. xiamenensis* NUITM-VS2 has been deposited at GenBank/EMBL/DDBJ under accession nos. AP026732 (chromosome), AP026733 (pNUITM-VS2_1), AP026734 [pNUITM-VS2_2 cocarrying *tet*(X4) and *tmexC3.2D3.2-toprJ1*], AP026735 (pNUITM-VS2_3), AP026736 (pNUITM-VS2_4 carrying *bla*_NDM-1_), and AP026737 (pNUITM-VS2 5).

## Ethical approval

Not required.

